# Subcortical nuclei of the human ascending arousal system encode anticipated reward but do not predict subsequent memory

**DOI:** 10.1101/2024.11.12.623212

**Authors:** Beth Lloyd, Steven Miletić, Pierre-Louis Bazin, Scott Isherwood, Desmond H. Y. Tse, Asta Håberg, Birte Forstmann, Sander Nieuwenhuis

## Abstract

Subcortical nuclei of the ascending arousal system play an important role in regulating brain and cognition. However, functional MRI of these nuclei in humans involves unique challenges due to their size and location deep within the brain. Here, we used ultra-high-field MRI and other methodological advances to investigate the activity of six subcortical nuclei during reward anticipation and memory encoding: the locus coeruleus, basal forebrain, median and dorsal raphe nuclei, substantia nigra and ventral tegmental area. Participants performed a monetary incentive delay task, which successfully induced a state of reward anticipation, and a 24-hour delayed surprise memory test. Region-of-interest analyses revealed that activity in all subcortical nuclei increased in anticipation of potential rewards as opposed to neutral outcomes. In contrast, activity in none of the nuclei predicted memory performance 24 hours later. These findings provide new insights into the cognitive functions that are supported by the human ascending arousal system.

## Introduction

Neuromodulation, governed by the subcortical ascending arousal system (AAS), plays a crucial role in regulating brain function and cognition. For example, dysfunction within the subcortex is widely implicated in the pathophysiology of various neuropsychiatric disorders (Shepherd 2013). However, despite its significance, the human AAS has remained relatively uncharted due to challenges in imaging small subcortical nuclei (Forstmann et al. 2017). Indeed, while animal models have provided valuable insights into the role of the AAS in cognition, human neuroimaging studies have often faced methodological limitations. Recent advancements in ultra-high field MRI offer a solution to some of these challenges, enabling more precise interrogation of neuromodulatory nuclei with improved spatial specificity and contrast-to-noise ratio (Hahn et al. 2013; Baecke et al. 2015; Priovoulos et al. 2018). Moreover, previous human imaging studies have primarily focused on the dopaminergic and noradrenergic systems, overlooking other key components of the AAS, such as the cholinergic and serotonergic systems. To our knowledge, only a handful of neuroimaging studies have examined multiple AAS nuclei in parallel (de Gee et al. 2017; Carvalheiro and Philiastides 2023; Lloyd et al. 2023; Mazancieux et al. 2023; Meissner et al. 2024), limiting a full understanding of the human AAS on cognition. In the present study, we utilized 7 Tesla (T) fMRI to examine the activity of six AAS nuclei as participants performed a monetary incentive delay task and a 24-hour delayed surprise memory test, providing the first comprehensive study on the role of the human AAS in two important cognitive functions: reward anticipation and memory encoding.

Previous 3 T fMRI studies have demonstrated reward anticipation effects in the dopaminergic midbrain (Wittmann et al. 2005; Schott et al. 2008; Wittmann et al. 2008). However, with the notable exception of Krebs et al (Krebs et al. 2011), most of these studies were unable to distinguish between the ventral tegmental area (VTA) and substantia nigra (SN). Animal studies have shown that AAS nuclei involved in the noradrenergic, cholinergic, and serotonergic systems are also phasically activated by reward-associated (i.e., conditioned) stimuli or tonically activated during the anticipation of reward delivery (Bouret and Sara 2004; Inaba et al. 2013; Nunez-Parra et al. 2020). Nevertheless, evidence of reward anticipation effects in human neuromodulatory nuclei *beyond* the dopaminergic midbrain remains scarce. Interestingly, both human (Schneider et al. 2018; Lloyd and Nieuwenhuis 2024) and non-human primate (Rudebeck et al. 2014) studies have reported that reward anticipation is characterized by a sustained increase in pupil size, a physiological marker that has been shown to reflect broader AAS activity (Joshi and Gold 2020). Therefore, it is likely that reward anticipation—an arousal-inducing phase of reward processing—engages neuromodulatory systems beyond the dopaminergic midbrain, also in humans.

In addition to examining reward anticipation effects, our study design allowed us to analyse the fMRI data collected during (incidental) encoding by categorising trials based on whether the items were later remembered or forgotten. This approach, known as the subsequent memory paradigm, has been instrumental in identifying regions within the prefrontal cortex and medial temporal lobe—particularly the hippocampus—where neural activity during encoding predicts later memory performance (Wagner et al. 1999). A smaller number of studies have paired this paradigm with pupillometry to explore peripheral indicators of AAS activity (for review, see Goldinger and Papesh 2012). Some of these studies have found subsequent memory effects reflected in pupil dilation, suggesting that phasic arousal may enhance memory encoding. Additionally, a small number of human neuroimaging studies have investigated the role of *specific* AAS nuclei in memory encoding, notably focusing on reward-related memories involving the dopaminergic VTA/SN (Wittmann et al. 2005; Gieske and Sommer 2023), as well as emotional memories linked to the noradrenergic locus coeruleus (LC) (Jacobs et al. 2020), providing preliminary insights into how specific neuromodulatory systems support memory encoding. Taken together, human pupillometry and neuroimaging research suggests that the AAS, as a whole, may play a broader role in memory formation beyond the dopaminergic and noradrenergic pathways, possibly due to its capacity to modulate arousal and attention, which are critical for prioritising information during encoding.

In the current study, we investigated whether reward anticipation and memory encoding were associated with neural activity in subcortical nuclei of the human AAS. We did not only differentiate between the VTA and SN but also examined BOLD responses in nuclei governing the noradrenergic (LC), cholinergic (basal forebrain, BF), and serotonergic systems (dorsal raphe nucleus, DRN; median raphe nucleus, MRN). Studying these small nuclei with fMRI presents unique challenges due to their size and location deep within the brain (Forstmann et al. 2017; Matt et al. 2019). To address these challenges, we implemented several dedicated methods. First, to ensure good signal quality, we used 7 T fMRI and a ‘tailored’ functional sequence optimized for imaging the subcortex (Miletić et al. 2020). Second, we derived high-resolution subcortical masks and used unsmoothed fMRI data, allowing for high spatial specificity (Bazin et al. 2020). Third, to suppress respiratory and cardiac artifacts, we recorded participants’ respiration and heart rate and applied a rigorous noise regression model (Glover et al. 2000; Harvey et al. 2008).

Thirty-two participants performed a monetary incentive delay task inside the 7T scanner, followed by a surprise memory test 24 hours later. During the monetary incentive delay task (**Figure 1A**), participants viewed man-made and natural items, which served as cues predicting whether a subsequent number classification task was rewarded or not (Wittmann et al. 2005). On each trial, participants were asked to first indicate whether they expected a reward (e.g., after man-made items) or not (after natural items), and then quickly classify a subsequent number as higher or lower than five. The difficulty of this number classification task was adapted so that roughly 70% of the reward trials were followed by a correct response and a reward, and the rest were followed by a small punishment. In the surprise memory test, participants were shown a mix of studied and unstudied items and were asked to classify each as ‘old’ or ‘new’, while also indicating their confidence in the response. This design allowed us to assess both recognition memory and confidence levels, and to examine how reward anticipation influenced memory performance. Behavioural analyses confirmed that the task successfully induced a state of reward anticipation, which subsequently enhanced memory performance. Our region-of-interest (ROI) analyses revealed that all AAS nuclei encoded reward anticipation. In contrast, while hippocampal subfields CA1 and CA3—areas that are known to be critical for the encoding of long-term memories (Frey et al. 1990)— predicted memory performance 24 hours later, none of the AAS nuclei showed a subsequent memory effect. These findings demonstrate that the opportunity to gain a reward mobilizes all neuromodulatory nuclei of the human AAS.

**Figure 1.**
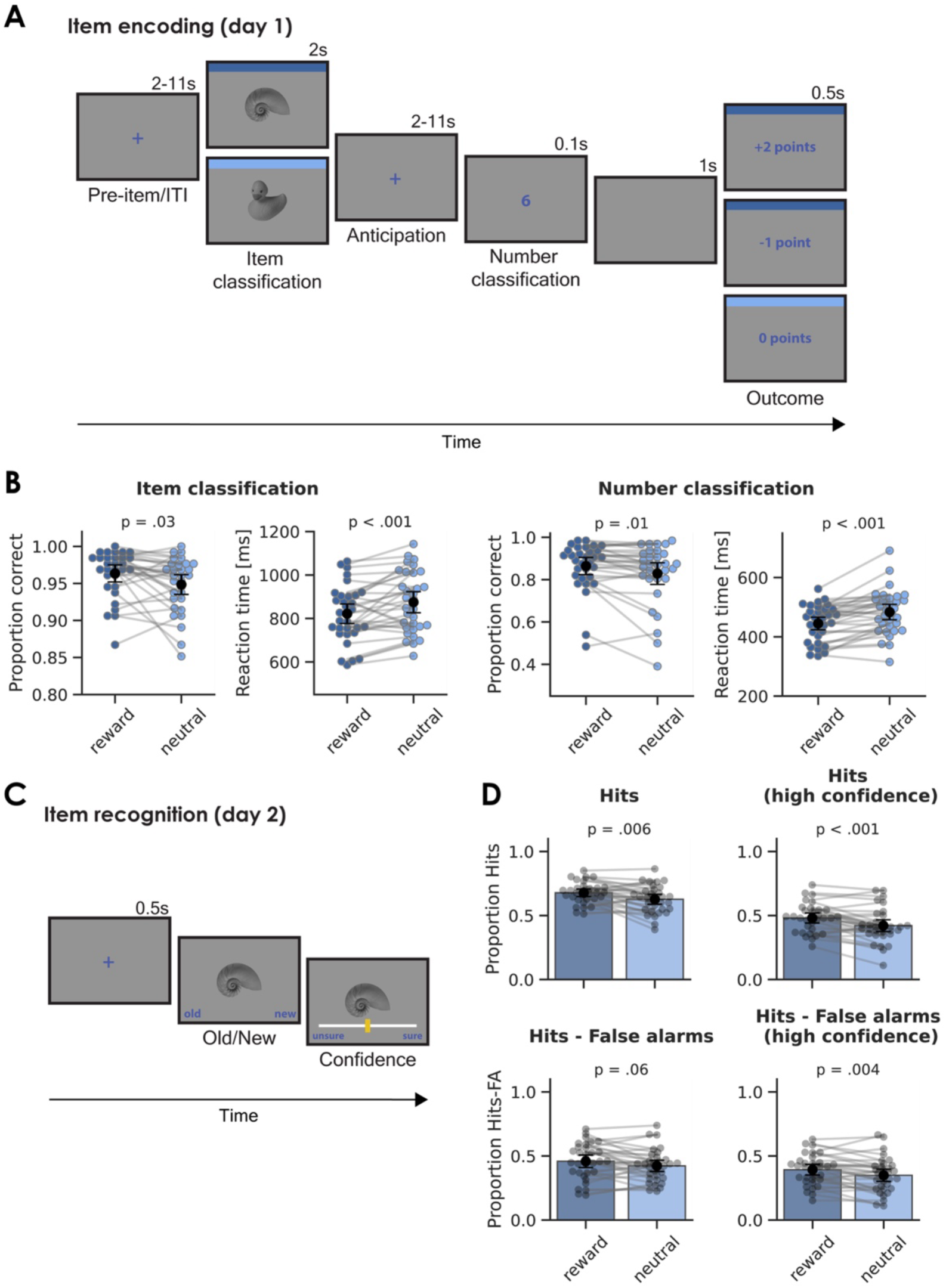
Manipulation successfully induced reward anticipation and reward anticipation boosted subsequent memory. A) Schematic overview of a trial in the monetary incentive delay task. Participants viewed images of man-made or natural items, which served as cues predicting whether a subsequent number classification task would be rewarded. Participants indicated whether they expected a reward (e.g., following man-made items; illustrated by the dark blue border) or not (e.g., following natural items; light blue border) before classifying a number as higher or lower than five. The difficulty of the classification task was adjusted so that approximately 70% of reward trials resulted in correct responses and rewards, while the remainder led to a small punishment. B) Response accuracy and reaction time on the item classification (left) and number classification task (right). C) A surprise memory test was administered 24 hours later, where participants judged a mix of studied and unstudied items as ‘old’ or ‘new’ and rated their confidence. D) Memory performance with hits (top) and recognition memory (hits – false alarms, bottom) separated by condition and for trials with high confidence. Data points and grey lines refer to individual participant scores. Black points refer to the group average and error bars indicate 95% confidence intervals.

## Materials and Methods

### Participants

A total of 32 participants took part in the current study (20 female; mean age 27.3 ± 6.2; age range 19-39). All participants were recruited from the Norwegian University of Science and Technology and were fluent in English, had normal or corrected-to-normal vision, and no history of epilepsy or overt clinical neuropsychiatric disease. Participants provided written informed consent and the experiment was approved by the Ethics Review Board of the University of Amsterdam (reference: 2021-BC-13146), and the Regional Committees for Medical and Health Research Ethics, Norway (reference: 116630). Two additional participants were excluded because they did not complete the memory test.

### Monetary incentive delay task

#### Stimuli

We used 384 man-made and natural items gathered from an openly available stimulus set (Brady et al. 2008) and Google Image Search. All items were grey-scaled and matched in luminance to the grey background (RGB: 125, 125, 125). Items were then resized (maximum width: 400 px, maximum height: 300 px) custom in-house scripts.

#### Design and procedure

The study consisted of two separate sessions: In session 1 participants carried out a monetary incentive delay task inside the scanner; in session 2, scheduled 24 h [± 2 h] later, they performed a surprise memory test. The monetary incentive delay task (**Figure 1A**) was programmed in Python using Expyriment (Krause and Lindemann 2014) and Psychopy2 (Peirce, 2007), and consisted of four runs of 64 trials each (256 trials in total). The trial sequence began with a fixation cross (duration 2-11 s, exponentially distributed with a mean of 5 s), followed by the presentation of a man-made or natural item (duration 2 s). Items were presented in a randomised order. Participants were told they could earn a reward during trials featuring items from one of two categories (“reward trials”), but not during trials with items from the other category (“neutral trials”). For half of the participants, man-made items (e.g., table, kettle, car) indicated a possible reward, while for the other half, natural items (e.g., flower, cat, shell) indicated a possible reward. During item presentation, participants were instructed to answer the question ‘Do you expect a reward on this trial?’ (item classification). To respond ‘yes’, participants pressed the right (green) button on the button box. To respond ‘no’, participants pressed the left (red) button on the button box.

Next, a fixation cross was presented (duration 2-11 s, pseudo-exponentially distributed with a mean of 5 s), followed by a target number (1, 4, 6, or 9 [randomised]; duration 0.1 s). After the target number appeared, participants were to respond as fast as possible to the question ‘Is the number higher or lower than 5?’ (number classification). To respond ‘higher’, participants pressed the right (green) button on the button box. To respond ‘lower’, participants pressed the left (red) button on the button box. The trial sequence ended with a blank screen (1 s), followed by the outcome for that trial (0.5 s). On reward trials, the outcome was either positive (‘+2 points’) or negative (‘-1 point’). On neutral trials, the outcome was always neutral (‘0 points’). Participants only received a positive outcome on reward trials if they responded correctly to the item classification and number classification questions and if their response to the number classification question met a response deadline. The response deadline was individually titrated based on the participant’s reaction times in all previous trials from that run to ensure that ∼70% of the reward trials ended with a positive outcome. Reward trials with any incorrect response or slow response to the number classification question ended with a negative outcome (**Figure 1A**).

After each run, participants were shown the number of points they earned. At the start of the session, participants were told they could earn points for good performance, and that the points would be converted into money. All participants then received 100 Kroner (more than the maximum possible earnings) at the end of the experiment. Before entering the MRI scanner, participants completed a practice of the monetary incentive delay task (∼ 5 minutes) using the same button box as they would use inside the scanner. Participants were instructed to press the buttons using their right index or middle finger.

### Surprise memory test

The next day, participants connected with the experimenter via a video call to complete the surprise memory test (**Figure 1C**), which was administered through Pavlovia (https://pavlovia.org/). The experimenter remained online (with the microphone on) while participants completed the memory test. Participants were initially informed they would be completing a similar task as on the previous day. The memory test was self-paced and consisted of 384 trials: the 256 items from the previous day intermixed with 128 lures. Each trial started with an item in the centre of the screen and underneath the item were the words ‘old’ (bottom left) and ‘new’ (bottom right). Participants indicated whether they had seen the item one day earlier (‘old’) or whether the item was completely new (‘new’). They made this response using the left and right arrow keys. Next, the item remained on the screen while a slider appeared underneath with the words ‘unsure’ (left side) and ‘sure’ (right side). Participants used the mouse to indicate how confident they were in their old/new response using the continuous slider from 0 (‘unsure’) to 100 (‘sure’). The memory test lasted ∼30 minutes.

### Image acquisition

MR data was acquired on a Siemens MAGNETOM TERRA (field strength = 7 T; gradient strength = 80 mT/m and maximum gradient slew rate = 200 T/m/s) using a 32-channel head coil. To collect anatomical data, we acquired a multi-echo gradient-recalled echo scan (GRE; TR = 31.0 ms, TE1 = 2.51 ms, TE2 = 7.22 ms, TE3 = 14.44 ms, TE4 = 23.23 ms, FA = 12°, field of view = 240 × 240 × 168 mm) and an MP2RAGE scan (TR = 4300 ms; TE = 1.99 ms; inversions TI1 = 840 ms, TI2 = 3270 ms; flip angle 1 = 5°, flip angle 2 = 6°, field of view = 240 ×240 ×168 mm; bandwidth = 250 Hz/Px; Marques et al., 2010). To collect BOLD activity data during the monetary incentive delay task, we acquired four functional echo-planar imaging (EPI) runs, each followed by four EPI volumes with opposite phase encoding direction for correcting susceptibility induced distortions. Functional images were recorded using a single-echo 2D-EPI BOLD sequence (TR = 1380 ms; TE = 14.0 ms; MB = 2; GRAPPA = 3; voxel size = 1.5 mm isotropic; partial Fourier = 6/8; flip angle = 60°; MS mode = interleaved; FOV = 192 ×192 ×128 mm; matrix size = 128 ×128; bandwidth = 1446 Hz/Px; slices = 82; phase encoding direction = A >> P; echo spacing = 0.8 ms). We also acquired a 2-average magnetization transfer-weighted turbo flash (MT-TFL) structural scan for delineation of the LC (TR = 400 ms; TE = 2.55; partial Fourier = 6/8; flip angle = 8°; voxel size = 0.4 x 0.4 x 0.5 mm; slices = 60) (Priovoulos et al. 2018) with the field of view placed perpendicular to the pons.

### Behavioural analyses

Trials were classified according to signal detection theory as either hits, misses, false alarms, or correct rejections. We assessed the proportion of remembered items (hits), as well as recognition memory, defined as proportions hits minus false alarms. For analyses that focused specifically on high-confidence responses, we selected trials where confidence was in the top third of the scale (> 66 out of 100). Any individual who scored more than 3 standard deviations away from the condition mean was treated as an outlier and removed from the relevant analysis. This outlier detection analysis resulted in the following exclusions: three participants on the basis of item classification accuracy, two participants on the basis of number classification accuracy and one participant on the basis of the proportion of hits. Note that these participants were included in the imaging analyses, but the key statistical results were identical if they were removed. Lastly, only trials with correctly classified items were used for the analyses involving memory performance.

### fMRI data preprocessing

All preprocessing of acquired anatomical and functional MRI data was carried out using *fMRIPrep* (Esteban et al. 2019; Esteban et al. 2020). Before registration, the structural (T1-weighted) image was corrected for intensity nonuniformity using *N4BiasFieldCorrection* (ANTs version 2.3.3) and skull-stripped with *antsBrainExtraction* using the OASIS30ANTs target template. For each of the four task runs, a BOLD reference volume and its skull-stripped version were generated. A fieldmap was estimated based on two EPI references with opposing phase-encoding directions using *3dQwarp* (AFNI). Based on this fieldmap, susceptibility distortion correction was applied to the EPIs, followed by applying boundary-based registration to the T1-weighted reference image using *bbregister* (FreeSurfer) with 6 degrees of freedom. Note that no fieldmap was acquired for one run from one participant, so fieldmap-free distortion correction was applied for this run. The BOLD reference volume was used to estimate 6 head-motion parameters (rotation and translation) with MCFLIRT (FSL version 5.0.9). Each of the four BOLD runs was then slice-time corrected to half of the acquisition range (0.674 s) using *3dT-shift* (AFNI).

The BOLD timeseries was denoised by applying an extensive noise correction model (41 confounds): i) 6 head motion estimates; ii) framewise displacement and DVARS (the spatial standard deviation of difference images), which were calculated for each functional run, both using their implementations in *Nipype*; iii) 12 discrete-cosine transform basis functions to model low-frequency scanner drifts; iv) to estimate the effects of physiological noise on the fMRI data, we recorded heart and breathing rate concurrently during fMRI data acquisition. We fit 18 regressors with retrospective image-based correction (RETROICOR) (Glover et al. 2000). PhysIO toolbox (Kasper et al. 2017) within the TAPAS software suite (Frässle et al. 2021) was used to estimate these physiological noise regressors. Note that physiological data was missing for all runs from one participant and for one run from another participant. Instead, the same number of *fMRIPrep*’s anatomic component-based noise correction (*aCompCor*) (Behzadi et al. 2007) regressors were entered in the model; v) we included one additional regressor to remove signal fluctuations from the fourth ventricle (mask computed using MASSP, details below) (Bazin et al. 2020). The fourth ventricle signal timeseries was mean-centered before it was added as a nuisance regressor.

### Definition of regions of interest

Seven subcortical ROIs were defined, six of which are part of the AAS: the LC, BF, MRN, DRN, SN, and VTA (**Figure 2A**). One, the striatum, was used as a control region for assessing reward anticipation effects. Aside from the LC, participant-specific probabilistic masks were computed for all subcortical ROIs (and the fourth ventricle) using the MASSP algorithm (Bazin et al. 2020). Three hippocampal subfields were selected as control regions for assessing subsequent memory effects. The CA1, CA3 and dendate gyrus (DG) were segmented based on individual T1-weighted images using *FreeSurfer* version 6.0 (*segmentHA_T1.sh*, labels FS60) (Van Leemput et al. 2009). Note that for bilateral ROIs (BF, SN, VTA, CA1, CA3, DG), left and right masks were defined, but the results presented here were collapsed across hemispheres. For visualization purposes, we created group-level ROI masks by transforming all participant-level masks into MNI-space and computing the average. See **Table S3** for an overview of the ROI masks.

**Figure 2.**
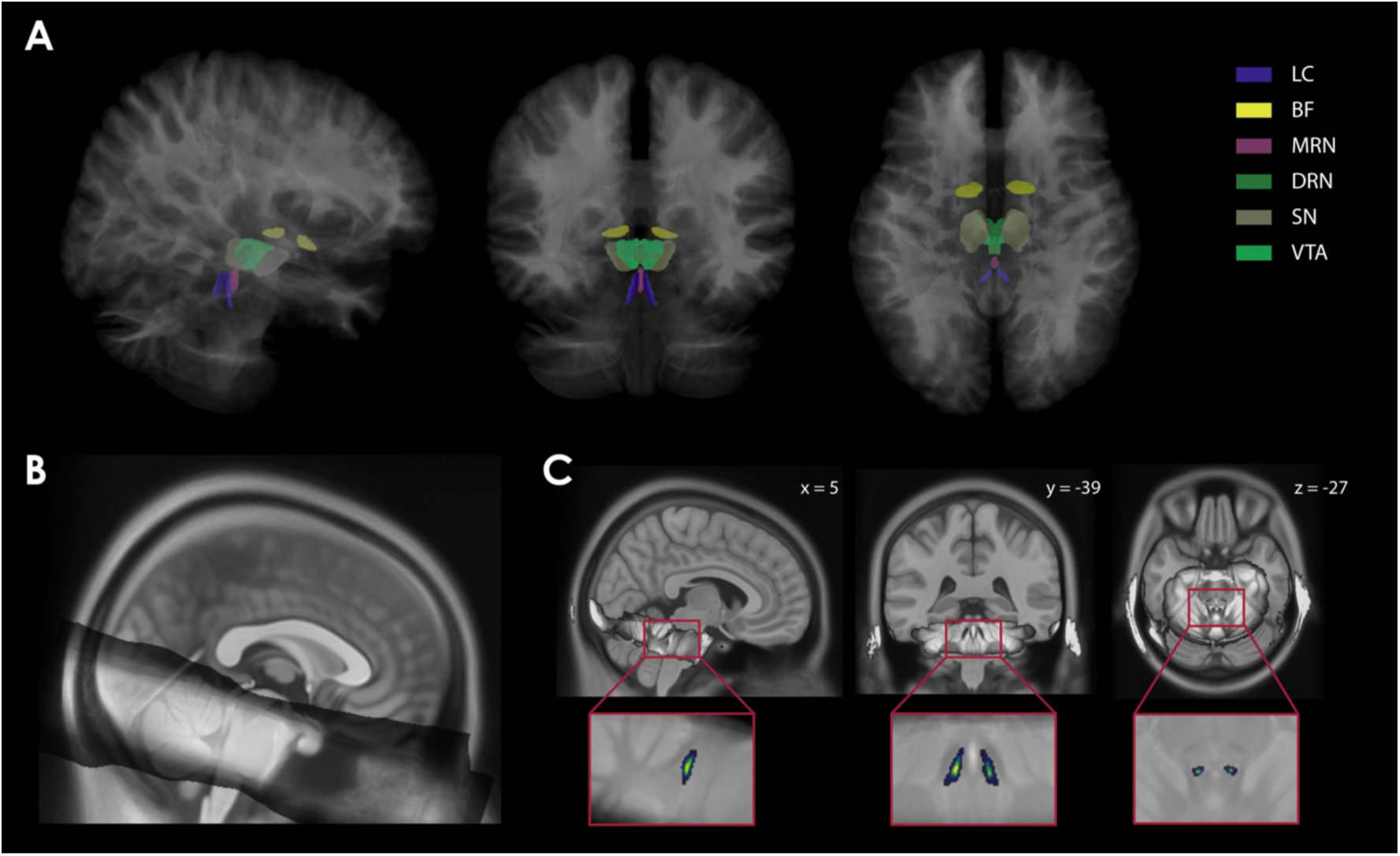
Overview of regions-of-interest (ROI) parcellation. A) Parcellations of AAS ROIs in MNI space (for visualization only). B) The group average MT-TFL image in MNI-space (grey), derived from the participant MNI-space MT-TFL images and overlaid on the MNI space template T1 image. C) The group-level probabilistic LC mask overlaid on the structural T1 (in MNI space for visualization only). LC, locus coeruleus; BF, basal forebrain; MRN, medial raphe nucleus; DRN, dorsal raphe nucleus; SN, substantia nigra; VTA, ventral tegmental area.

To define the LC mask, the following steps were performed in ANTs version 2.3.3 (Tustison et al. 2010). First, N4 bias field correction was applied to the two MT-TFL scans of each participant. Next, we made a participant-specific average MT-TFL image using *antsMultivariateTemplateConstruction2*, which resulted in a processed MT-TFL image. For each participant we co-registered the processed MT-TFL image with their unprocessed (with skull) T1-weighted scan. With the transformation matrices produced by this registration, and the linear and non-linear transforms acquired from warping the processed participant T1-weighted image to the MNI-space template, we moved the processed MT-TFL image from participant space into group (MNI) space (using *antsApplyTransforms*). Using these participant MNI-space MT-TFL images we computed a group average MT-TFL image (see **Figure 2B**). From this, we segmented the LC using a semi-automated segmentation approach utilizing a previously published 7 T probabilistic LC mask (Ye et al. 2021) as a starting point. To that end, we first estimated the weighted mean and standard deviation of intensity of the group average MT-TFL image within the previously published 7 T probabilistic LC mask. We used this as a lower cutoff of intensity, by transforming the average with a sigmoid centered on the mean and with the slope corresponding to the standard deviation. Next, we performed the same procedure, but this time we centered it at the mean plus three times the standard deviation (*µ + 3σ*) to define high intensities (i.e., nearby cerebral spinal fluid [CSF]). We dilated the published LC mask by 0.5 mm, and did the same for the CSF sigmoid map (but not the first sigmoid map). Finally, we multiplied the dilated LC mask, the first sigmoid, and 1 minus the CSF sigmoid, and took the square root to create our group-level probabilistic LC mask, combining the spatial information from the previous LC mask with the intensity information from the group average MT-TFL image (see **Figure 2C**). Lastly, the group-level probabilistic LC mask was transformed back into participant-specific T1-weighted space using the relevant transformation matrices, resulting in the final participant-level LC masks. Since the fourth ventricle and LC are in close proximity to each other, we ensured that no voxels were included in the LC mask that might belong to the fourth ventricle for each participant. To achieve this, we compared the participant-level LC and fourth ventricle probabilistic masks for overlapping voxels. When overlapping voxels were found, the voxel with the highest probability was retained in its respective mask, while it was removed from the other mask. Note that the group-level probabilistic LC mask was only used for visualization purposes.

### fMRI data analyses

We extracted the timeseries from each ROI using the computed probabilistic masks. These timeseries were extracted from unsmoothed functional images to ensure regional specificity. The contribution from each voxel in the mask to the mean signal of the region was weighted by its probability of belonging to that region. Timeseries were then converted to percentage signal change by dividing the timeseries by the mean signal value, multiplying by 100, and subtracting 100. Next, task events were convolved with a canonical double gamma hemodynamic response function. These included our event of interest (item onset, duration 2 s) as well as the rest of the events in a trial (number onset and outcome onset, duration 0.001 s). To identify contrasts for reward anticipation and subsequent memory effects, trials were modelled with four different regressors based on condition (reward, neutral) and memory (remembered, forgotten). The general linear model (GLM) design matrix also contained the noise correction model described above (41 confounds). The model was fit using ordinary least squares. The estimated BOLD responses (beta coefficients) were calculated per run, per ROI for each participant. These betas were then used as a dependent variable in higher-level analyses. ROI-based GLMs were performed using package *nideconv* (De Hollander et al. 2019).

In addition to this GLM, we performed two additional GLMs: one assessing the level of confidence as a parametric modulator for memory effects (GLM 2) and one to assess the reward effects while controlling for reaction time duration on the item classification task (GLM 3). For GLM 2, confidence ratings on the surprise memory test were z-scored and added as a modulator in the model. For GLM 3, reaction time duration on the item classification task was modelled as a separate regressor (Mumford et al. 2024). This was treated both as a regressor of interest and as a confound in our control analysis.

To corroborate the ROI-based reward anticipation effects observed in the current study, we performed voxel-wise GLM analyses on the whole brain, specifically to inspect whether the whole-brain BOLD contrast (reward vs. neutral trials) is in line with the literature. Prior to fitting this whole-brain GLM, functional data were spatially smoothed using a SUSAN (Smith and Brady 1997) (kernel size full width half maximum = 4.5 mm). Task regressors were convolved with the canonical HRF and first-level GLMs were fit with the same design matrix and noise correction model as for the ROI analysis. Whole-brain analyses were run using the FILM method from FSL FEAT. In a second-level GLM, we used fixed effects analyses to combine the resulting run-level GLMs. We then estimated group-level models using FLAME1 and FLAME2 from FSL. SPMs were generated to visualize the resulting group-level contrast. The maps were corrected for the false discovery rate (FDR) using a critical value of q < 0.05 (Yekutieli and Benjamini 1999).

For one participant, only two runs of data were acquired (they wanted to exit the scanner) and one participant had no positive feedback in one run, so that run was removed. In addition, runs with > 50% incorrectly classified trials were removed from all analyses (nine runs across five participants). Finally, using an additional nuisance regressor, we removed trials with items that were incorrectly classified. This regressor (2 s) was added to the design matrix for the specific run (included in ROI-based and whole-brain GLMs).

### Statistical analyses

Statistical analyses were carried out in Python 3 (frequentist) and Rstudio version 4.0.3 (Bayesian). To estimate the potential effects of reward anticipation on item and number classification and subsequent memory, we conducted paired-samples *t*-tests (α ≤ 0.05). To test for reward anticipation effects and subsequent memory effects on fMRI BOLD signal in our ROIs, we fit linear mixed-effects (LME) models using the *mixedlm* function from the statsmodels package. In these models, the item-related betas from the ROI-based GLMs were predicted by condition (reward, neutral) and subsequent memory (remembered, forgotten). To account for individual deviations from fixed group effects, intercepts for participants were modeled as random effects. The resulting *p*-values for the ten ROIs were FDR-corrected. In addition, to quantify the evidence for any effects of interest, we ran Bayesian LME models using *stan_lmer* from the rstanarm package in R. Throughout, data are expressed as the mean ± standard error of the mean (SEM).

## Results

This section is organized as follows: We first verified the success of the reward manipulation through examining the behavioural data from the monetary incentive delay task. Next, we evaluated memory performance to determine if rewarded items were better remembered than neutral items. Finally, we determined the effects of reward anticipation and memory encoding success on the activity of our AAS nuclei, as well as a number of control regions.

### Behavioural data

#### Manipulation successfully induced reward anticipation

During the monetary incentive delay task (**Figure 1A**), participants responded to two questions, both related to the reward anticipation manipulation. First, they were presented with a cue, and responded to the question ‘Do you expect a reward on this trial?’ (item classification). Second, they were presented with a digit, and they responded to the question ‘Is the number higher or lower than 5?’ (number classification). The effects of the task manipulation on reaction time and accuracy of these responses are shown in **Figure 1B**. Overall, participants performed well on the item classification task, but they showed higher accuracy on reward trials (96 ± 3% SE) than on neutral trials (95 ± 4 %; *t*(29) = 2.3; *p* = .03). Reaction times were also significantly shorter on reward trials (822 ± 130 ms) than on neutral trials (875 ± 140 ms; *t*(31) = -3.53; *p* < .001), although this was not necessary to attain a reward. These results suggest that the reward cue (item) invigorated participants, prompting them to perform better in anticipation of the potential reward. A similar pattern was found for the number classification task: accuracy was higher (reward: 86 ± 12%; neutral: 83 ± 15%; *t*(30) = 2.7; p = .01) and reaction times were faster (reward: 445 ± 61 ms; neutral: 483 ± 75 ms; *t*(31) = -4.92; *p* < .001) on reward trials, on which task performance determined the outcome of the trial, than on neutral trials.

#### Reward anticipation enhanced memory

Previous studies have shown that reward anticipation strengthens memory encoding (Wittmann et al. 2008; Wittmann et al. 2011). Consistent with these findings, our study demonstrated that reward anticipation effectively boosted memory formation, as tested after a 24-h delay. The results from the surprise memory test (**Figure 1C**) are presented in **Figure 1D**. The proportion of hits was significantly higher for reward-associated items (0.68 ± 0.08) than for items with no reward association (0.63 ± 0.11; *t*(30) = 2.96; *p* = .006). This pattern was also observed for high-confidence hits (reward: 0.48 ± 0.11, neutral: 0.42 ± 0.13; *t*(31) = 4.32; *p* < .001). In addition to assessing the proportion of hits, we tested whether recognition memory (hits minus false alarms) differed between conditions. Recognition memory was better for reward items (0.39 ± 0.12) compared to neutral items (0.35 ± 0.14; *t*(31) = 3.1; *p* = .004) when only high-confidence trials were considered, but not when all trials were included (reward: 0.46 ± 0.14, neutral: 0.42 ± 0.13; *t*(31) = 96; *p* = .06). These findings align with previous research in showing beneficial effects of reward anticipation on specific aspects of memory performance.

### fMRI data

To estimate the item-related BOLD response for reward and neutral trials, we fit a GLM to the timeseries extracted from individually-derived probabilistic masks for each ROI in subject specific space (**Figure 2A**). A weighted mean timeseries was computed for each ROI, with weights based on the probability of each voxel belonging to the region. We then used ROI-specific linear mixed-effects (LME) models to predict the GLM BOLD response estimates, with condition (reward, neutral) and subsequent memory (remembered, forgotten) as predictors (see *Materials and Methods*). To quantify the relative strength of the evidence for these effects, we carried out Bayesian model comparisons between a condition-only model, a memory-only model, a model with condition and memory as joint predictors, and an intercept-only model. The models were compared using Bayes factors, which quantify the relative likelihood of the models (Kass and Raftery 1995; Ly et al. 2016).

#### Reward anticipation broadly increased AAS activation

First, we tested for a reward anticipation effect in a control region, the striatum, where reward-related BOLD responses are expected (Miller et al. 2014). As anticipated, the estimated BOLD response in the striatum was significantly larger for reward-associated items than for neutral items (*z* = 6.48, *pcorr* < .001). Next, we inspected the reward anticipation contrast in the BOLD data from our AAS ROIs. As shown in **Figure 3**, we found a significant effect of reward anticipation in all AAS ROIs: LC (*pcorr* = .004), BF (*pcorr* = .006), MRN (*pcorr* < .001), DRN (*pcorr* = .046), SN (*pcorr* < .001), and VTA (*pcorr* < .001). These results are important because, until now, only the SN and VTA have been shown to encode anticipated reward in humans. Our findings highlight the broader involvement of the AAS in reward processing.

**Figure 3.**
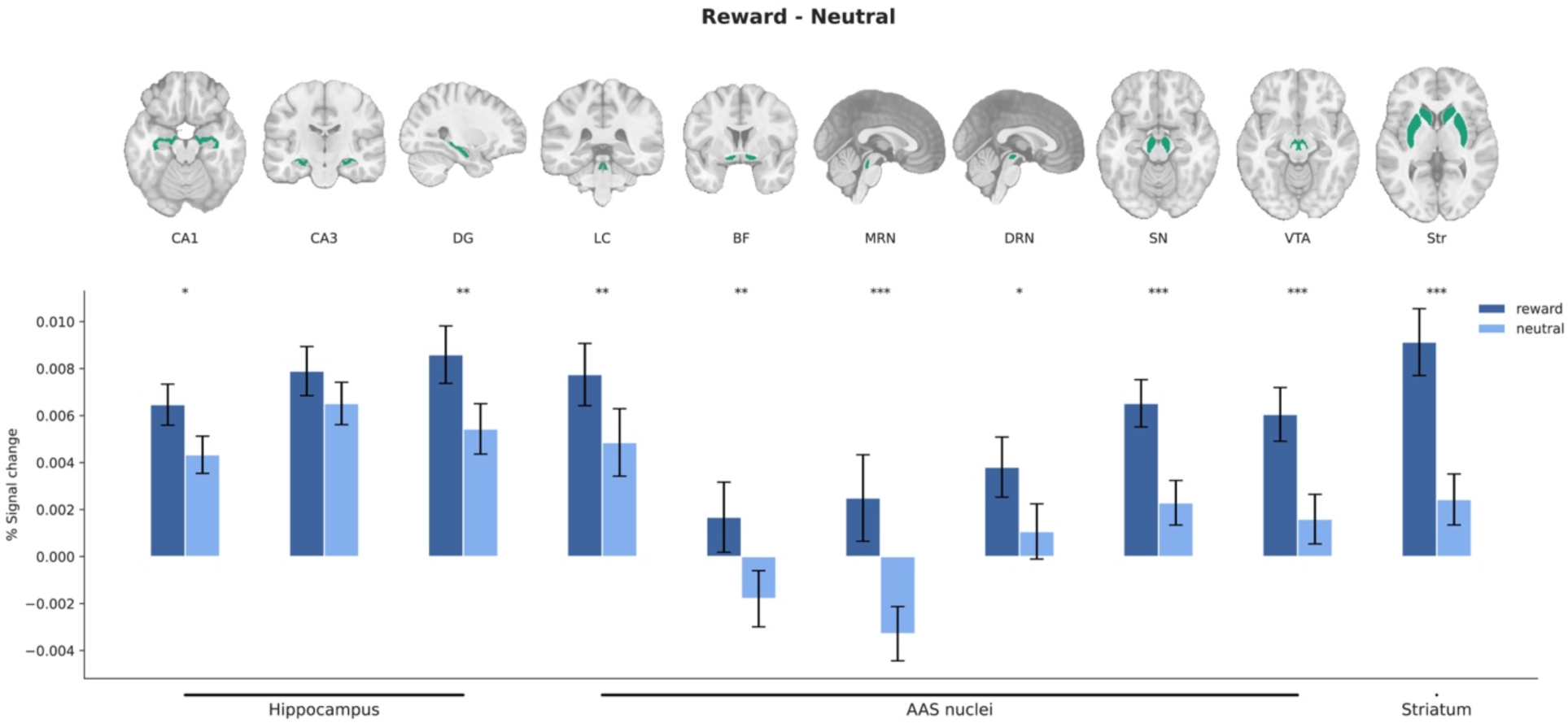
Ascending arousal system (AAS) nuclei encode anticipated reward. Masks are shown for each ROI (green; in group space for visualisation only). Bar plots illustrate reward anticipation effects. Significant reward anticipation effects were observed in all AAS ROIs, as well as in the hippocampal subfields CA1, DG, and Str (*p* < .05: *; *p* < .01: **; *p* < .001: ***; FDR-corrected for all ROIs). Error bars represent 95% confidence intervals. DG, dendate gyrus; LC, locus coeruleus; BF, basal forebrain; MRN, medial raphe nucleus; DRN, dorsal raphe nucleus; SN, substantia nigra; VTA, ventral tegmental area; Str, Striatum.

Reward anticipation effects were also present in two of the hippocampal ROIs (CA1 and DG; all *pcorr* < .015), and in some other subcortical and cortical regions (**Figure S1**).

#### Hippocampal subfield but not AAS activation was associated with stronger memory encoding

Next, we investigated whether the BOLD response in any AAS ROIs exhibited a subsequent memory effect. Before doing this, we verified whether later remembered items were associated with increased BOLD responses in the hippocampus. Based on previous literature, we expected these effects to occur in hippocampal subfields CA1, CA3, and DG, which are frequently associated with encoding of long-term memories (Frey et al. 1990; Frey et al. 1991; Huang and Kandel 1995; Sariñana et al. 2014). Employing the same LME models used to quantify reward anticipation effects, we found that BOLD response estimates in subfields CA1 and CA3 were significantly larger for items later remembered than for items later forgotten (CA1: *z* = 3.36, *pcorr* = .01, CA3: *z* = 2.63, *pcorr* = .04; **Figure 4**), with no effect in the DG (*z* = 1.9, *pcorr* = .20).

**Figure 4.**
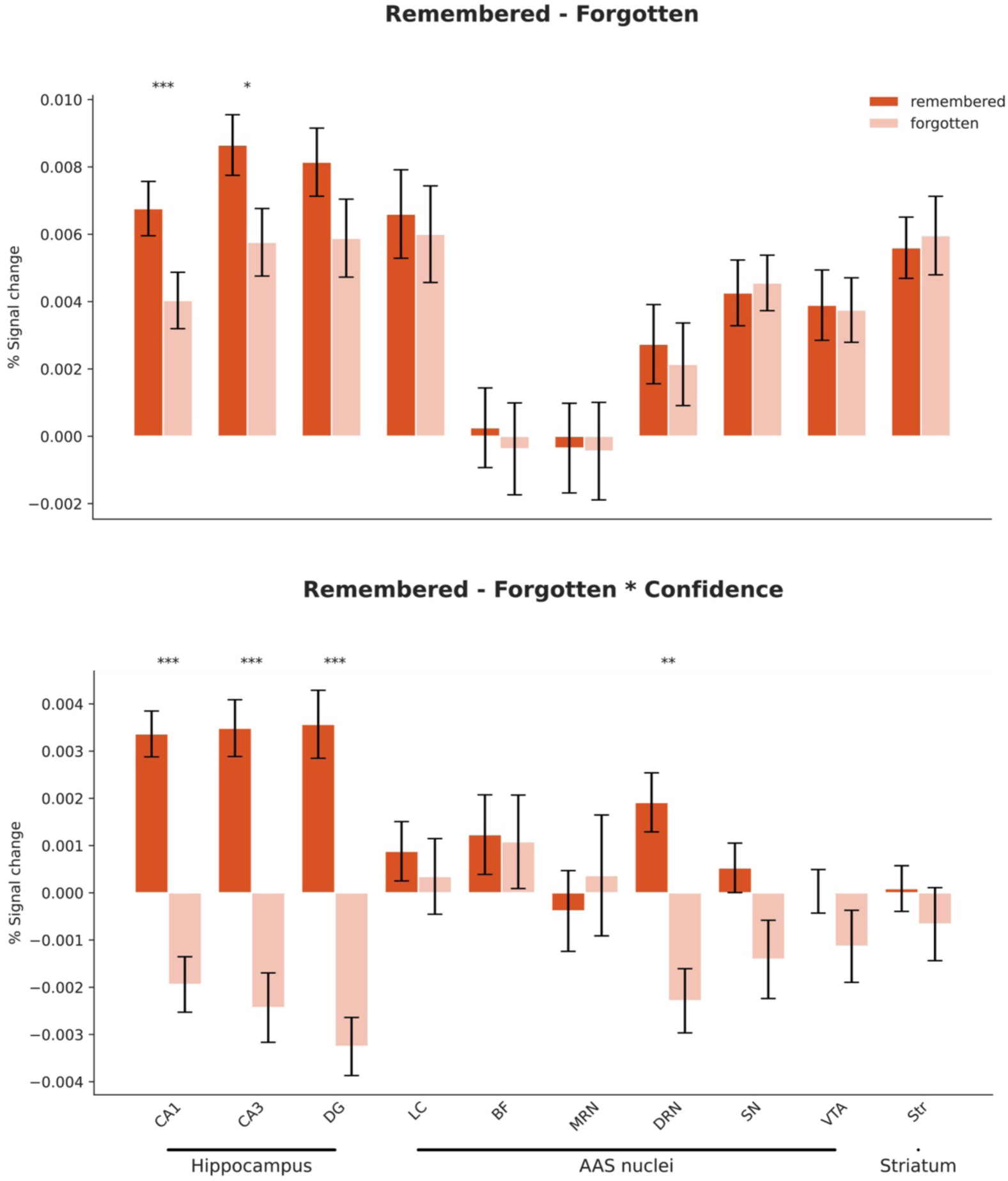
Subsequent memory effects are largely limited to the hippocampal subfields. Bar plots illustrate subsequent memory effects. Significant effects were observed in the hippocampal subfields CA1 and CA3, as well as in the DRN and DG (the latter only when BOLD responses were modulated by confidence level). No subsequent memory effects were detected in any of the AAS ROIs (*p* < .05: *; *p* < .01: **; *p* < .001: ***; FDR-corrected for all ROIs). Error bars represent 95% confidence intervals. DG, dendate gyrus; LC, locus coeruleus; BF, basal forebrain; MRN, medial raphe nucleus; DRN, dorsal raphe nucleus; SN, substantia nigra; VTA, ventral tegmental area; Str, Striatum.

Interestingly, we found no evidence for a subsequent memory effect in any of the AAS nuclei (all *p*corr > .75; **Figure 4**). Indeed, Bayesian model comparisons showed that the preferred overall model for each AAS ROI (except the DRN) was the model that included only condition as a predictor. Specifically, for most ROIs we found moderate to very strong evidence in favour of the condition-only model, as indicated by the following Bayes factors: LC (BF10 = 3.59), BF (BF10 = 2.55), MRN (BF10 = 159.98), DRN (BF10 = 0.92), SN (BF10 = 478.31), and VTA (BF10 = 373.10). For a detailed overview of the Bayesian model comparisons, refer to **Table S1**.

To investigate how the BOLD response during item encoding varied as a function of confidence ratings for each item, we repeated the GLM but now incorporating confidence as a parametric modulator. We were specifically interested in whether confidence ratings modulated the BOLD signal in our AAS nuclei for remembered vs. forgotten items. Again, we first inspected the hippocampal subfields and observed significantly higher BOLD signal estimates in the CA1, CA3 and DG for remembered items compared to forgotten items when confidence ratings were included as a parametric modulator (CA1: *z* = 7.1, *pcorr* < .001, CA3: *z* = 7.1, *pcorr* < .001, DG: *z* = 7.4, *pcorr* < .001; **Figure 4**). No effect was found in any AAS nuclei (all *p*corr> .25) except the DRN, which showed larger activation for later remembered items (*z* = 3.2, *p*corr= .002), mirroring the hippocampal subfields. Note that the beta weights for several ROIs were negative for forgotten items (**Figure 4** bottom). This is intuitive when one considers that the correlation between confidence and BOLD response differs in sign between remembered and forgotten items; remembered items show a positive relationship, while forgotten items exhibit a negative relationship between confidence and BOLD response in the AAS.

#### Activation of four AAS nuclei increases with reaction time

Recent pupillometry studies (Murphy et al. 2016; Gross and Dobbins 2021) have suggested a role for the AAS in generating decision urgency, an evidence-independent signal that expedites the evolving decision process by driving it closer to a fixed decision threshold (Carland et al. 2019). Because urgency mounts over the course of the item presentation duration until a decision is made, AAS activity should be positively correlated with reaction time (RT), a prediction that is borne out by pupil-dilation data (Murphy et al. 2016; Gross and Dobbins 2021). Here, we examined if any of the AAS nuclei showed a positive trial-by-trial relationship between BOLD activation and RT in the item classification task. To this end, we reran the original GLM while including RT as an additional regressor with varying durations. Interestingly, one-sample *t*-tests revealed that the regression weights of the RT predictor showed a significant positive association with BOLD responses in four AAS nuclei: the LC (*p*corr < .001), MRN (*p*corr < .001), DRN (*p*corr = .003), and VTA (*p*corr < .001) (**Figure 5**). Although the nature of these results is correlational, these results are in line with a causal role for the AAS in generating decision urgency. Note that these positive BOLD-RT relationships cannot account for our reward anticipation effects on AAS BOLD responses, because reward anticipation produced a *negative* relationship between BOLD response (larger) and RT (shorter). Indeed, as shown in **Table S2**, reward anticipation effects persisted in all our AAS ROIs after we controlled for RT duration (Mumford et al. 2024).

**Figure 5.**
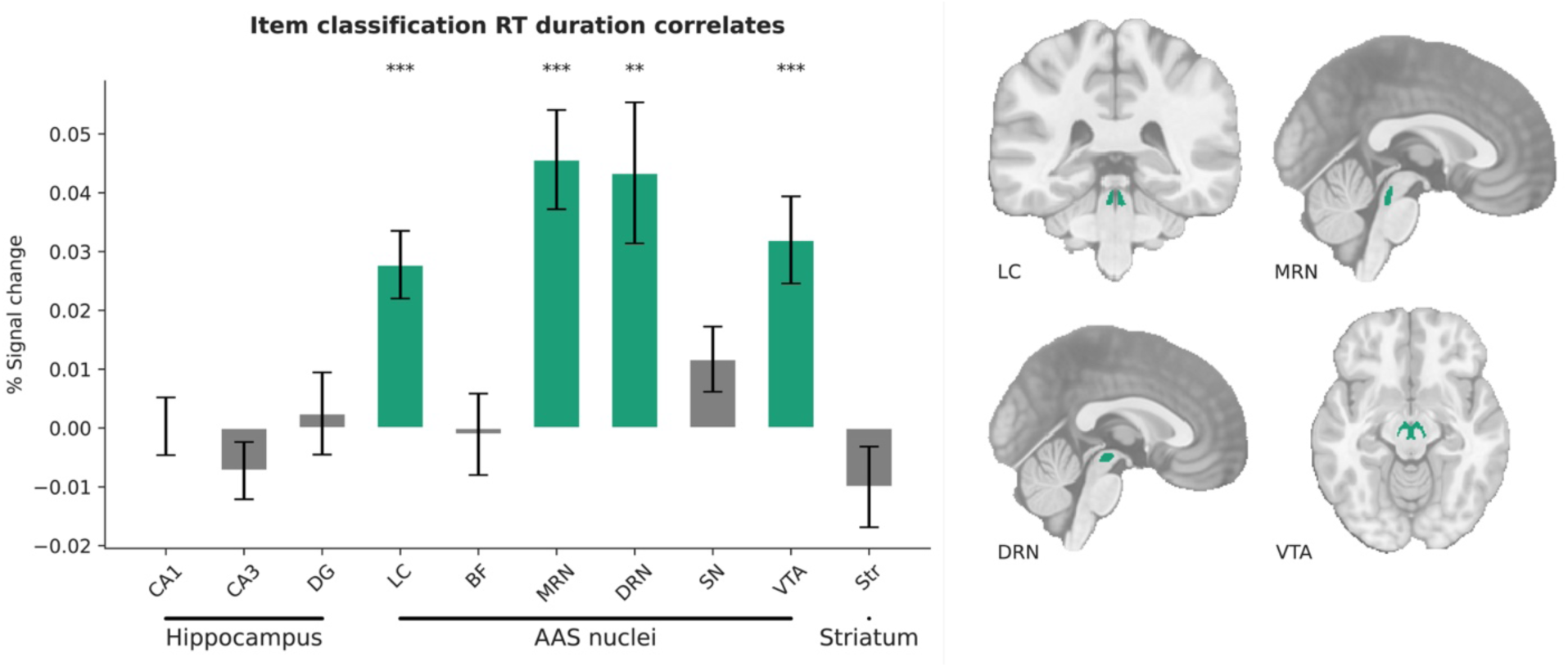
Reaction time on the item classification response task predicts activation of specific Ascending arousal system (AAS) nuclei. Green bars indicate ROIs where BOLD activation and RT were positively correlated across trials. A significant correlation between RT duration and BOLD signal was observed in the LC, MRN, DRN, and VTA (*p* < .05: *; *p* < .01: **; *p* < .001: ***; FDR-corrected for all ROIs). Masks from these regions are visualised on the structural T1 image. Error bars indicate 95% confidence intervals. DG, dendate gyrus; LC, locus coeruleus; BF, basal forebrain; MRN, medial raphe nucleus; DRN, dorsal raphe nucleus; SN, substantia nigra; VTA, ventral tegmental area; Str, Striatum.

## Discussion

In this study, we employed ultra-high field 7 T fMRI to investigate the activity of six subcortical nuclei within the AAS (LC, BF, MRN, DRN, SN, VTA) during reward anticipation and memory encoding. In the first session, participants completed a monetary incentive delay task while undergoing fMRI. Consistent with previous studies using similar rewarded reaction time tasks (Wittmann et al. 2005; Adcock et al. 2006; Wittmann et al. 2011; Lloyd and Nieuwenhuis 2024), our task successfully induced a state of reward anticipation, evident through faster and more accurate behavioural responses when a reward was at stake. The following day, participants completed a surprise memory test. Again, in line with previous findings (Wittmann et al. 2005; Wittmann et al. 2008; Wittmann et al. 2011; Murayama and Kitagami 2014), we observed memory benefits for items that were previously predictive of a reward compared to neutral items. Critically, using high-resolution fMRI data and individually-derived probabilistic masks, we demonstrated that activity in all AAS nuclei increased in anticipation of potential rewards as opposed to neutral outcomes. In contrast, unlike two of the hippocampal subfields that we examined, none of the AAS nuclei predicted subsequent memory. These findings provide new insights into the cognitive functions that are supported by the AAS.

Previous human fMRI studies have found reward anticipation effects in the dopaminergic midbrain (Wittmann et al. 2005; Schott et al. 2008; Wittmann et al. 2008). However, with one notable exception (Krebs et al. 2011), these studies have generally been unable to anatomically distinguish between the VTA and SN. In this study, we not only demonstrate distinct reward anticipation effects in the VTA and SN but also show increased activation in the LC, BF, MRN, and DRN in response to reward-related items compared to neutral ones. These effects remained after we controlled for trial-to-trial fluctuations in RT duration, which can significantly confound estimated BOLD responses (Mumford et al. 2024). Our findings align with a body of research conducted on the AAS in rodents and non-human primates. It is well established that dopamine neurons in the VTA and SN exhibit phasic firing in response to reward-predicting cues (Hollerman and Schultz 1998; Cohen et al. 2012). However, although less well documented, the AAS nuclei governing the noradrenergic, cholinergic and serotonergic systems are also phasically activated by reward-associated (i.e., conditioned) stimuli and/or tonically activated during the anticipation of reward delivery (Bouret and Sara 2004; Inaba et al. 2013; Nunez-Parra et al. 2020). Together, these findings suggest that all neuromodulatory nuclei of the AAS are responsive to the prospect of future rewards. Our task design does not allow us to distinguish whether the phasic neural responses underlying these reward anticipation effects reflect the subjective value of the anticipated reward, a signed or unsigned reward prediction error (i.e., discrepancy between expected and actual rewards), or some related quantity (Feng et al. 2024; Jordan 2024). This remains an intriguing question for future research.

In addition to BOLD modulations, we found clear behavioural effects of reward anticipation. First, reward anticipation was associated with improved performance (higher response speed and accuracy) on the item and number classification tasks. This is consistent with previous findings that feature-based attention, as defined by the rate of information uptake, is enhanced for reward-predicting stimuli compared to neutral stimuli (Spaniol et al. 2011; Dix and Li 2020). The incentive salience hypothesis (Berridge and Robinson 1998) proposes that these positive effects are mediated by reward-related dopamine signals that strengthen the perceptual salience assigned to the neural representations of reward-associated items, thereby enhancing attention to these items. Second, reward-associated items were remembered better than neutral items. Reward-related memory enhancements are known to be mediated by interactions between midbrain dopamine regions and the hippocampus (Shohamy and Adcock 2010). In line with this literature, we found that reward anticipation modulated BOLD responses not only in the VTA and SN but also in hippocampal subregions CA1 and DG.

In addition to the reward-neutral contrast, we also analysed the fMRI data from the encoding phase by categorizing trials based on whether items were later remembered or forgotten. Previous studies have combined similar subsequent memory paradigms with pupillometry to assess peripheral markers of AAS activity during encoding. Some of these studies found that larger pupil size during encoding was positively associated with later memory success (reviewed in Goldinger and Papesh 2012), suggesting that phasic arousal may strengthen memory encoding. Based on these findings, we anticipated that at least some AAS nuclei would exhibit a similar subsequent memory effect in our data. However, although all AAS nuclei were engaged during reward anticipation, none predicted memory performance 24 hours later. Of course, potential subsequent memory effects in AAS nuclei may be more subtle than reward anticipation effects. Deep brain nuclei are generally smaller and have weaker signal-to-noise ratio than cortical regions, making subtle effects harder to detect. However, we did detect subsequent memory effects in the hippocampal subregions CA1 and CA3, which had masks similar in size to the VTA and SN, and signal-to-noise ratios comparable to those of the LC and MRN (**Table S3**). Furthermore, our Bayesian model comparisons yielded moderate to strong evidence against a subsequent memory effect in the AAS nuclei. In conclusion, an association between AAS BOLD activity and subsequent memory is unlikely.

These null results seem inconsistent with two previous human fMRI studies that found a subsequent memory effect in the VTA/SN (Wittmann et al. 2005; Gieske and Sommer 2023). We consider three possible explanations. First, Wittman (Wittmann et al. 2005) observed subsequent memory effects in the VTA/SN after a retention interval of three weeks, as compared to 24 hours in our study. The relatively short retention interval in our study may have contributed to the absence of subsequent memory effects in the AAS (Wittmann et al. 2005; Düzel et al. 2010). However, this is difficult to reconcile with the reported subsequent memory effects in Gieske and Sommer (Gieske and Sommer 2023), whose recognition test took place “the next day” (p. 4529). Second, we assessed memory success using an old-new recognition test, and focused our fMRI analysis on a comparison between all ‘hits’ and all ‘misses’. In contrast, the two previous studies distinguished between recollection and familiarity, analysing only recollected items (those judged as ‘remembered’) but not familiar items (those judged as ‘known’). This raises the question whether VTA/SN activity during encoding may specifically affect recollection rather than familiarity. A third distinction between the three studies is that Wittman (Wittmann et al. 2005) and Gieske and Sommer (Gieske and Sommer 2023) searched for individual voxels in the VTA/SN that survived a particular statistical threshold (i.e. “peak voxels”), while we averaged across all voxels in each AAS ROI. Because BOLD fMRI is affected by various types of noise, selected peak voxels may include voxels in which the noise happened to correlate with the subsequent memory regressor, potentially resulting in false positives. On the other hand, our averaging approach makes it harder to detect functionally specialized (e.g., memory-enhancing) subregions within a given ROI. In addition to these three possible explanations, other methodological differences, such as variations in voxel resolution, field strength, and spatial smoothing, may have introduced partial volume effects, further contributing to the discrepancies between the studies.

Dopamine and noradrenaline play an important role in the preferential encoding in the hippocampus of reward related and novel events (Shohamy and Adcock 2010; Takeuchi et al. 2016). However, we are not aware of any animal studies that have linked *trial-to-trial fluctuations* in AAS activity during encoding and the consequences of those fluctuations for later memory. Nor did we find any neurophysiological evidence that directly speaks to our finding that DRN activation showed an interaction between memory success and confidence level, ranging from lowest activation for ’old items categorized as new (i.e., misses) with high certainty’ to medium activation for ’old items categorized as old or new with low certainty’ to highest activation for ’old items categorized as old (i.e., hits) with high certainty’. Although in the memory test the categorical old/new judgment was separated from the requirement to indicate confidence level on a graded scale, the virtual scale ranging from ’sure new’ to ‘unsure’ to ’sure old’ is often interpreted as reflecting familiarity strength (Wixted et al. 2010). This raises the intriguing possibility that in our study, fluctuations in DRN activity during encoding contributed to variability in familiarity strength during the test phase. This possibility warrants further investigation in future studies.

There is growing evidence from pupillometry studies that the AAS generates decision urgency (Murphy et al. 2016; Steinemann et al. 2018; Gross and Dobbins 2021; Poth 2021; Lloyd and Nieuwenhuis 2024; Tromp et al. 2024), an evidence-independent signal that rises over the course of a decision process, driving evidence accumulators toward a fixed decision threshold for committing to an action (Carland et al. 2019). Agents are thought to “add” this dynamic urgency signal to all evidence accumulators when faced with (mild to large) time pressure, to maximize their subjective reward rate (Carland et al. 2019). Because urgency level rises over the course of a trial until a decision is made, any physiological marker of urgency should be positively related to trial-to-trial fluctuations in RT. Maximum pupil dilation shows this signature of dynamic urgency (Murphy et al. 2016; Gross and Dobbins 2021; Lloyd and Nieuwenhuis 2024), suggesting that phasic arousal generates dynamic urgency. Here, we found that BOLD activation in four AAS nuclei (LC, MRN, DRN, VTA) was also positively correlated with RT, regardless of whether the item predicted a reward or a neutral outcome. Consistent with a role in urgency generation, the firing rate of the monkey LC increases rapidly just before a speeded instrumental response, but not before a non-task-related response (Clayton et al. 2004; Bouret and Richmond 2009). Although RT-BOLD correlations have been regarded as confounding fMRI analyses (Yarkoni et al. 2009; Mumford et al. 2024), brain areas exhibiting such correlations may include areas that are the source or target of decision urgency.

Imaging subcortical nuclei poses several methodological challenges (Forstmann et al. 2017; Matt et al. 2019). Firstly, the small size of these nuclei makes them susceptible to partial-volume averaging at conventional spatial resolutions (de Hollander et al. 2015). To address this, we employed high-resolution functional imaging and avoided spatial smoothing in our fMRI data. Secondly, deep subcortical nuclei have unique magnetic properties, such as high iron concentrations, which complicate functional imaging (Posse et al. 1999). To optimize contrast in these iron-rich subcortical structures, we used a shorter echo time than is typical in conventional fMRI, so as to maximize BOLD contrast in the subcortex (Miletić et al. 2020). Thirdly, the proximity of the AAS nuclei to air-filled cavities, cerebrospinal fluid, and major arteries makes them particularly vulnerable to movement and other sources of physiological noise. To limit these artefacts, we implemented a rigorous noise regression model, accounting for measured cardiac and respiratory signals, as well as residual signal from the fourth ventricle. Fourthly, accurately delineating smaller subcortical structures is crucial for obtaining meaningful results, yet their localization can be challenging. ROI masks are often defined at the group level using published coordinates or probabilistic atlases. However, due to individual variability in subcortical structures, these masks may not be precisely accurate (Fjell et al. 2013; Keuken et al. 2014). We addressed this by creating individual masks for each participant using the multi-contrast anatomical subcortical structures parcellation (MASSP) algorithm (Bazin et al. 2020). Because the LC is notoriously difficult to precisely localize and register (Yi et al. 2023), we implemented a different set of dedicated, state-of-the-art methods: i) a specialised 7 T MRI sequence sensitive to LC contrast, enabling high-resolution imaging of the LC for each individual (Priovoulos et al. 2018), ii) ANTs SyN for optimized brainstem co-registration, as ANTs has been identified as the best-performing tool for normalization (Ewert et al. 2019), and iii) a semi-automated segmentation approach using predefined boundaries and surrounding CSF to create a customised probabilistic atlas for the LC based all individuals in the sample. A caveat of our imaging study is that the scans, although acquired at 7 T, do not capture the anatomical and functional heterogeneity of AAS nuclei. The DRN, for example, harbours a population of dopamine neurons that responds to rewarding (and aversive) events (Cho et al. 2017), and may have contributed to the reward anticipation BOLD responses reported here.

While a lot of research in cognitive neuroscience has focused on the functions of the *individual* systems in the AAS (e.g., the dopamine system), less attention has been given to how these systems interconnect. Evidence suggests that these neuromodulatory systems share notable similarities in their anatomical organization (Briand et al. 2007) and connectivity (España and Berridge 2006; Sara 2009; Gugel et al. 2024), and exhibit direct interactions with one another (Avery and Krichmar 2017). Despite this, studies exploring their co-activation and interactions remain relatively scarce. Besides the current study, only a handful of human fMRI studies have simultaneously examined multiple components of the AAS (de Gee et al. 2017;

Carvalheiro and Philiastides 2023; Lloyd et al. 2023; Mazancieux et al. 2023; Meissner et al. 2024). The AAS BOLD responses in the present study showed a remarkably uniform pattern of results: all nuclei showed a reward anticipation effect and none showed a subsequent memory effect (except the DRN in interaction with confidence level). Previous fMRI studies have also found broad co-activation of AAS nuclei in response to locally and globally deviant stimuli (Mazancieux et al. 2023) and in relation to pupil size (de Gee et al. 2017; Lloyd et al. 2023; Meissner et al. 2024). Together, these fMRI findings reinforce the idea that these systems are often recruited in concert, influence each other and simultaneously modulate brain and cognition (Briand et al. 2007).

## Acknowledgments

This work was supported by funding from the Vici grant awarded to Sander Nieuwenhuis, financed by the Dutch Research Council (NWO), with project number VI.C.181.032. We would also like to thank Sarah Habli, Lisbeth Røe, and Daniel R. Sokolowski for help with data collection.

## Author contributions

B. Lloyd: Conceptualization, Data curation, Formal analysis, Investigation, Methodology, Validation, Visualization, Writing –original draft. S. Miletić: Data curation, Formal analysis, Investigation, Methodology, Writing –review & editing. PL. Bazin: Methodology, Software, Visualization, Writing –review & editing. S.J.S. Isherwood: Data curation, Project administration, Writing –review & editing. D.H.Y. Tse: Data curation, Methodology, Project administration, Writing –review & editing. A.K. Håberg: Project administration, Resources, Writing –review & editing. B.U. Forstmann: Methodology, Project administration, Resources, Writing –review & editing. S. Nieuwenhuis: Conceptualization, Funding acquisition, Investigation, Methodology, Project administration, Resources, Supervision, Writing –review & editing.

## Data and code availability

Code used to preprocess and analyse the data for the current study can be found here: https://github.com/bethlloyd/subcortical_nuclei_of_the_human_AAS

The data acquired in this study will also be uploaded and freely available online (www.figshare.com) after full collection of the database and anonymization is possible.

Any additional information required to re-analyse the data reported in this paper is available from the corresponding author upon reasonable request.

