## Supplementary material for "Subcortical nuclei of the human ascending arousal system encode anticipated reward but do not predict subsequent memory": Figure S1

**Supplementary Materials**

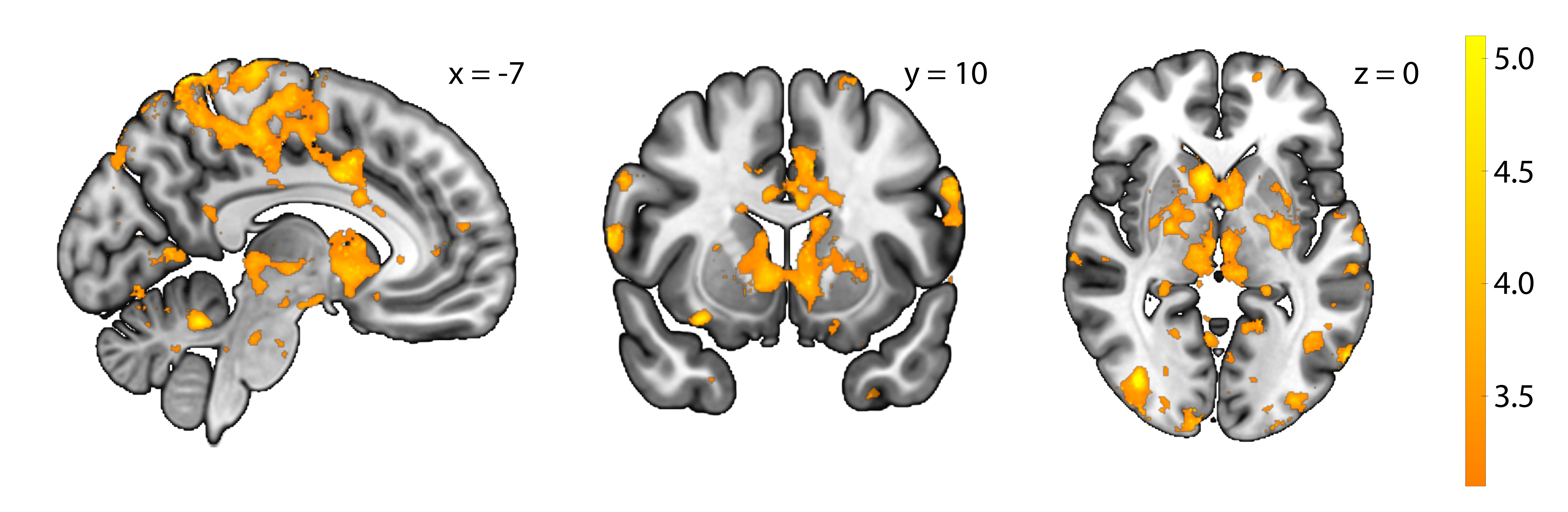

**Figure S1.** Group-level statistical parametric map for the reward anticipation contrast across the whole brain in MNI-space. Activation maps are thresholded using false discovery rate (FDR) correction (q < 0.05, z-value = 3.1).

**Table S1.**

|  | **Preferred Model** | **BF_10_** | **Evidence for alternative** |
| --- | --- | --- | --- |
| **CA1** | Reward + Memory | 18.42 | Strong evidence |
| **CA3** | Memory | 2.08 | Anecdotal evidence |
| **DG** | Reward | 4.03 | Moderate evidence |
| **LC** | Reward | 3.59 | Moderate evidence |
| **BF** | Reward | 2.55 | Anecdotal evidence |
| **MRN** | Reward | 159.98 | Very strong evidence |
| **DRN** | Intercept | 0.92 | Anecdotal evidence against |
| **SN** | Reward | 478.31 | Very strong evidence |
| **VTA** | Reward | 373.1 | Very strong evidence |
| **Str** | Reward | 5.8 x 10^6^ | Very strong evidence |

*Note.* Results of Bayesian model comparisons for each ROI. The preferred model refers to the model with the highest Bayes factor. The null model includes only the intercept, while the alternative models include a condition-only predictor, a memory-only predictor, or condition and memory as joint predictors. Bayes factors are presented as BF_10_, where values greater than 1 indicate evidence in favor of the alternative model (preferred model), and values lower than 1 indicate evidence in favor of the null model (intercept-only). DG, dendate gyrus; LC, locus coeruleus; BF, basal forebrain; MRN, medial raphe nucleus; DRN, dorsal raphe nucleus; SN, substantia nigra; VTA, ventral tegmental area; Str, Striatum. BF: Bayes Factor.

**Table S2.**

|  | **z** | ***p*** | ***p_corr_*** | **significance** |
| --- | --- | --- | --- | --- |
| **CA1** | 1.6 | 0.110 | 0.122 |  |
| **CA3** | 1.04 | 0.299 | 0.299 |  |
| **DG** | 1.93 | 0.054 | 0.067 |  |
| **LC** | 2.73 | 0.006 | 0.012 | * |
| **BF** | 2.33 | 0.020 | 0.033 | * |
| **MRN** | 3.62 | 0.000 | 0.000 | *** |
| **DRN** | 2.16 | 0.031 | 0.044 | * |
| **SN** | 3.42 | 0.001 | 0.002 | ** |
| **VTA** | 3.48 | 0.001 | 0.002 | ** |
| **Str** | 4.8 | 0.000 | 0.000 | *** |

*Note.* Statistics for the reward anticipation contrast after controlling for RT (*p* < .05: *; *p* < .01: **; *p* < .001: ***). *p*_corr_ refers to the *p*-values after correcting for FDR. DG, dendate gyrus; LC, locus coeruleus; BF, basal forebrain; MRN, medial raphe nucleus; DRN, dorsal raphe nucleus; SN, substantia nigra; VTA, ventral tegmental area; Str, Striatum.

**Table S3.**

|  | **mean tSNR** | **mean N voxels** | ***source*** |
| --- | --- | --- | --- |
| **CA1** | 40.81 | 292.96 | freesurfer |
| **CA3** | 43.34 | 103.62 | freesurfer |
| **DG** | 44.57 | 164.46 | freesurfer |
| **LC** | 41.54 | 59.69 | custom scripts |
| **BF** | 26.26 | 79.53 | MASSP |
| **MRN** | 40.39 | 37.5 | MASSP |
| **DRN** | 37.09 | 49.56 | MASSP |
| **SN** | 30.52 | 285.94 | MASSP |
| **VTA** | 36.84 | 164.07 | MASSP |
| **Str** | 49.7 | 3975.37 | MASSP |

*Note.* Mean tSNR refers to the transient signal-to-noise ratio, averaged across participant-level masks. Mean N voxels refers to the number of voxels in functional space (1.5 mm isotropic), averaged across all participant-level masks. Source: freesurfer (van Leemput et al., 2009); MASSP (Bazin et al., 2020); MT-TFL (magnetization transfer-weighted turbo flash; Priovoulos et al., 2018). DG, dendate gyrus; LC, locus coeruleus; BF, basal forebrain; MRN, medial raphe nucleus; DRN, dorsal raphe nucleus; SN, substantia nigra; VTA, ventral tegmental area; Str, Striatum.
